## Supplementary figures and images for "A *Drosophila* Model of Pontocerebellar Hypoplasia Reveals a Critical Role for the RNA Exosome in Neurons"

### Fig S1

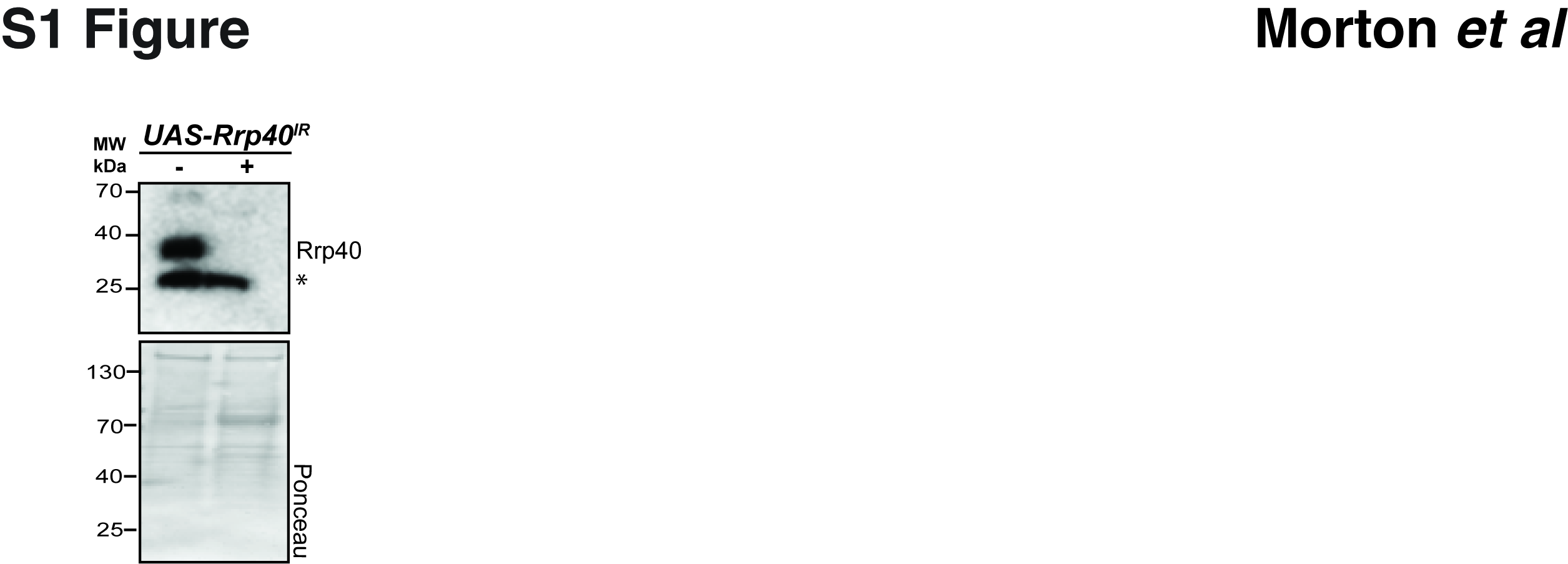

### Fig S2

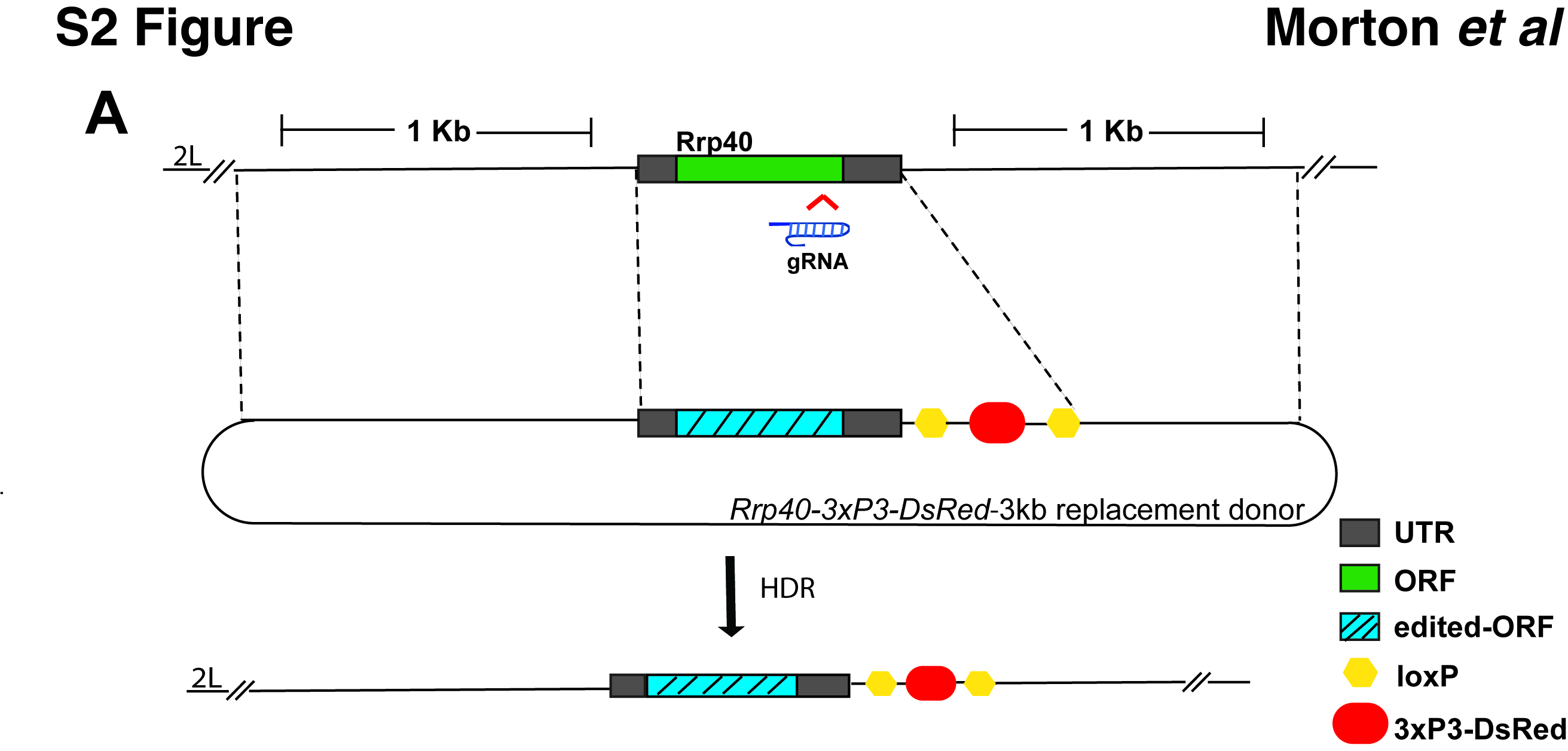

### Fig S3

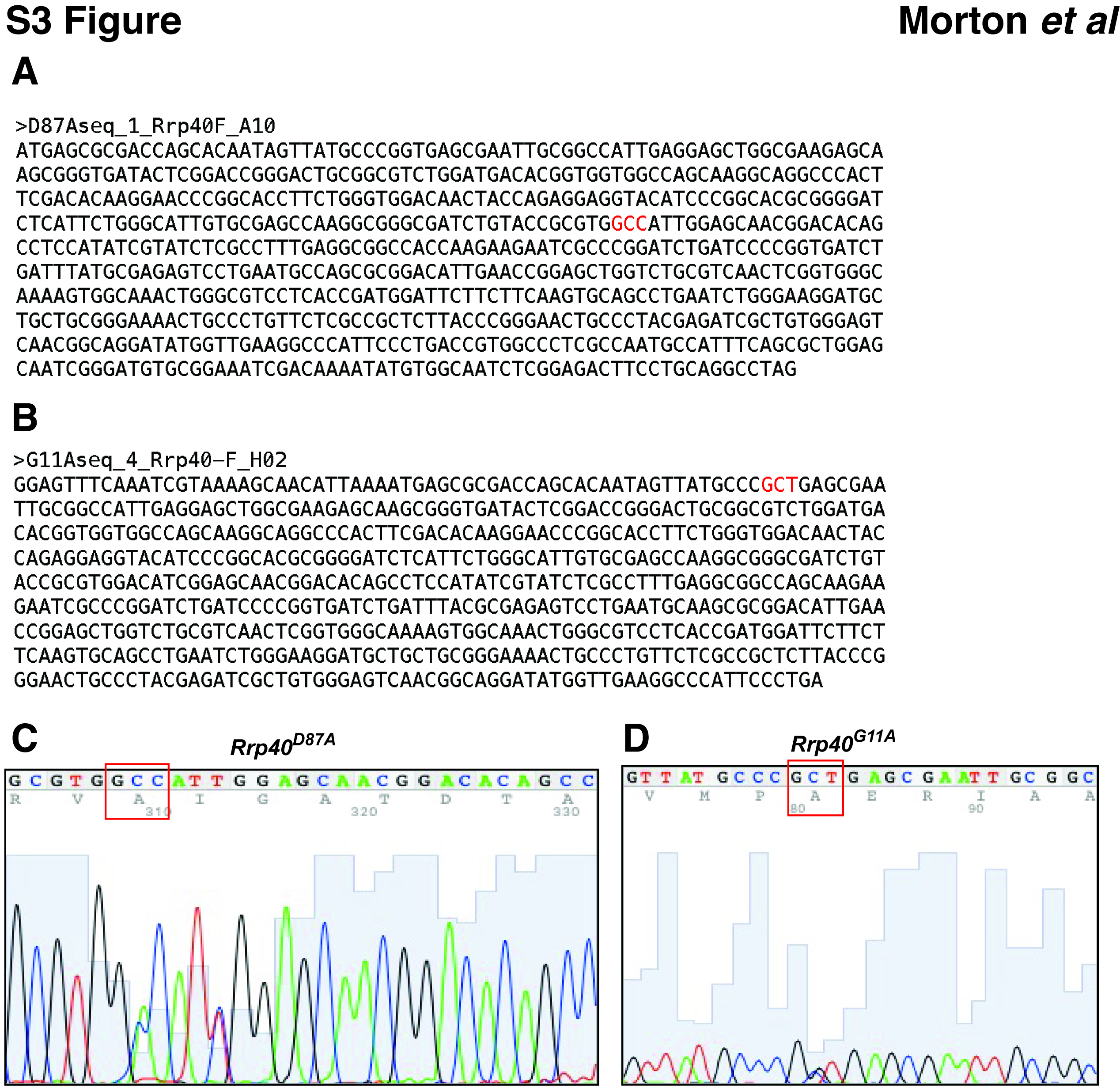

### Fig S4

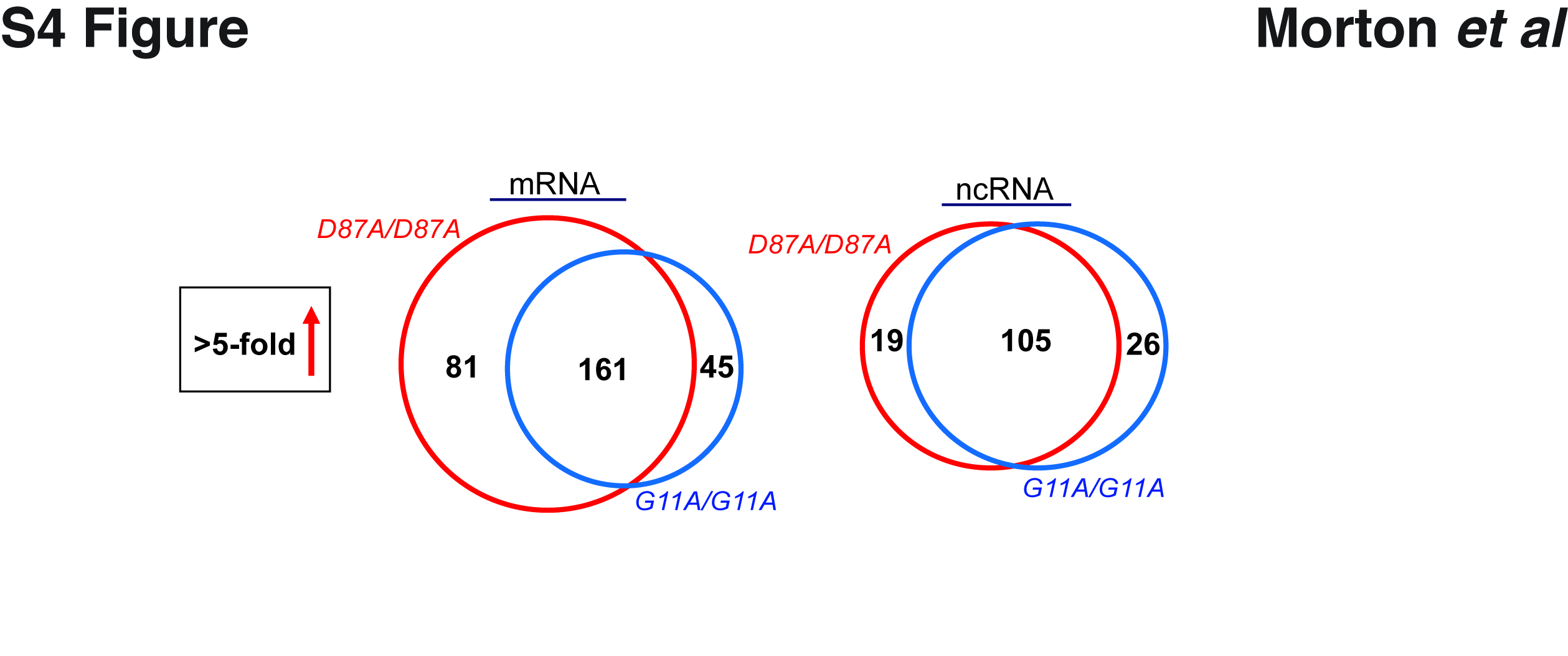

### Fig S5

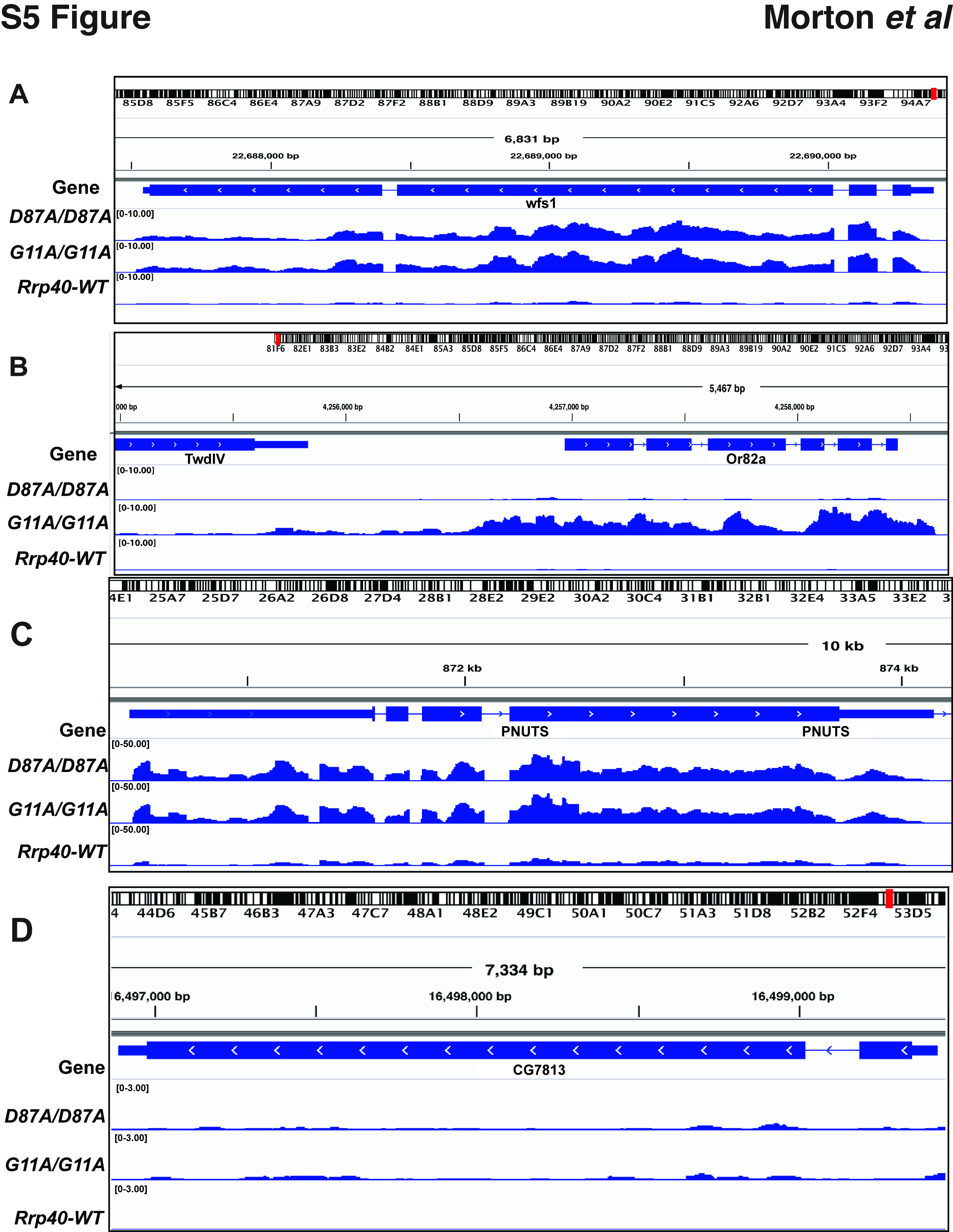

### Fig S6

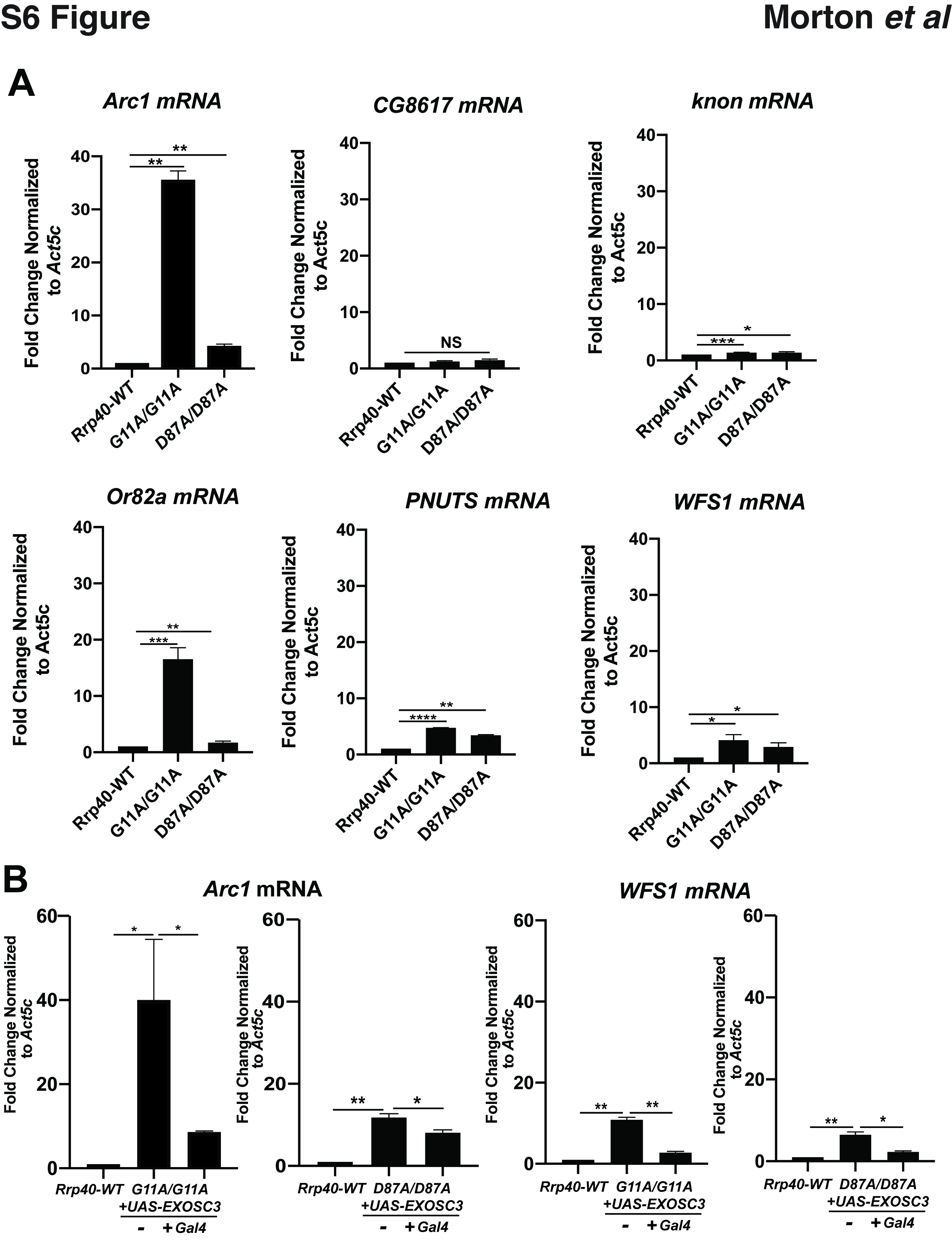
